# The landscape of structural variation in coppery titi monkeys (*Plecturocebus cupreus*)

**DOI:** 10.64898/2026.01.13.699302

**Authors:** Cyril J. Versoza, Karen L. Bales, Jeffrey D. Jensen, Susanne P. Pfeifer

## Abstract

Coppery titi monkeys (*Plecturocebus cupreus*) are an important non-human primate model for studying neurobiology and social behavior, in part owing to their relatively unusual combination of social monogamy and paternal care. Despite this importance, relatively little is known regarding the underlying population genomics of this platyrrhine. This study presents high-coverage, whole-genome sequencing data from 26 individuals which, combined with a highly accurate multi-algorithm ensemble approach, was used to characterize the first map of structural variation in the species. This novel genomic resource includes over 13,000 structural variants, with the majority (>90%) being copy number variants. While many of these were found to be located in intergenic regions, several affected genes associated with disease, including an inversion predicted to impact a pathway implicated in early-onset Parkinson’s disease. Furthermore, utilizing parent-offspring trios included within this study, the *de novo* structural variant rate was estimated to be one in every 1.5 births, similar to that reported in rhesus macaques but considerably higher than that observed in large human cohorts, as may be expected from underlying differences in life history traits amongst these species. Taken together, these insights into the structural variant landscape of *P. cupreus* will not only improve their utility as a behavioral model system, but will also contribute to our general understanding of the role of structural variation in both the evolution of the primate clade and disease-outcomes.

## INTRODUCTION

Copy number variants — genomic segments larger than 50 bp that vary in DNA dosage between individuals (including deletions and duplications) — and other structural variants that change genome organization (including insertions and inversions), are one of the largest sources of heritable variation (Conrad et al. 2010; 1000 Genomes Project Consortium 2015; Chaisson et al. 2019; Collins et al. 2020). As such, they play an important role in shaping population genetic and phenotypic variation within and between species (Hollox et al. 2022). In addition to potentially facilitating adaptation in novel and changing environments, the divergence of structural variants amongst populations can also impact the degree of gene flow, and thereby contribute to reproductive isolation and speciation (see the reviews of Iskow et al. 2012 and Kondrashov 2012, and the commentary by Feulner and De-Kayne 2017). Moreover, structural variants impacting coding and regulatory regions can have profound effects on health and disease, particularly with regards to the susceptibility to infectious agents (see the review of Hollox and Hoh 2014) as well as neurodevelopmental, neurological, and psychiatric disorders including autism spectrum disorder, schizophrenia, attention deficit hyperactivity disorder, Alzheimer’s and Parkinson’s disease (see the review of Harner et al. 2025 and references therein).

Historically studied using molecular cytogenic techniques (such as fluorescence in situ hybridization and comparative genomic hybridization) and chromosomal microarray technologies, the emergence of whole-genome sequencing technologies has recently provided a valuable and cost-effective alternative to identify copy number and other structural variants at the population-scale. For paired-end short-read data specifically — which remains the *de facto* standard in the field due to both lower costs and sample requirements relative to those necessary for long-read technologies — several structural variant callers have been developed that scan the genome for regions that exhibit variations in read coverage, harbor an excess of split read alignments, and/or read pairs with discordant insert sizes or conflicting strand orientation (for an overview of the different detection strategies, see Figure 2 in Alkan et al. 2011). In contrast to the more widely studied single nucleotide variation (Pfeifer 2017), standardized guidelines still need to be established for the identification of structural variation (Ho et al. 2020); however, while individual short-read structural variant callers can suffer from high false discovery rates, the application of multiple methodologies leveraging different (and often complementary) signals — referred to as an “ensemble” approach — has generally been shown to improve both sensitivity and specificity compared to single-caller approaches (see the benchmarking studies of Cameron et al. 2017; Kosugi et al. 2019; Gabrielaite et al. 2021; Kosugi and Terao 2024). Indeed, recent research has demonstrated that, while long-read data is necessary to comprehensively characterize structural variants in repetitive regions of the genome, short-read data exhibit a similar level of precision and recall for different types of structural variants in non-repetitive regions as well as for large (> 1 kb) deletions in repetitive regions (see Figure 4 in Kosugi and Terao 2024).

Due to these computational and technological advances, recent years have witnessed a renewed interest in the study of structural variation in many organisms. In primates, for example, differences in copy number variation have been observed between the genomes of anthropoid apes and other haplorrhines, contributing to the divergence between species (e.g., Gokcumen et al. 2013; Sudmant et al. 2013; Dennis et al. 2017; Kronenberg et al. 2018; Li et al. 2020b). In two of the most comprehensive studies of structural variation in primates to date, Porubsky et al. (2020) used single-cell and long-read sequencing to catalogue inversions across the great apes, whereas Mao et al. (2024) combined information from genome assemblies of eight non-human primates (chimpanzee, bonobo, gorilla, orangutan, gibbon, macaque, owl monkey, and marmoset) together with short- and long-read sequencing data to catalogue deletions and insertions fixed in the human lineage, showing that >25% of the genome has been impacted by structural variation. While long-read sequencing data remains limited for many species, population-specific catalogs of structural variation have also been generated from short-read data for several organisms. For instance, Brasó-Vives et al. (2020) and Thomas et al. (2021) constructed genome-wide maps of copy number variation from 198 and 32 rhesus macaques — the most frequently used non-human primate model in biomedical research — finding that, similar to humans, a large proportion of the species’ genome is impacted by structural variation. Furthermore, comparing the genomes of 60 rhesus macaques from five Chinese populations and one Indian population, Liu et al. (2025) identified several population-specific copy number variants suggested to be related to environmental adaptation and phenotypic variability (such as body size). Similarly, sequencing the genomes of 38 common marmosets — another species of biomedical relevance — Yang et al. (2023) discovered a deletion enriched in individuals displaying an epileptic phenotype. More recently, based on short-read data from 14 individuals, Versoza et al. (2025) provided the first insights into the structural variation characterizing the genome of the aye-aye (and see the review of Soni et al. 2025) — representing one of the most basal splits in the primate tree. Yet, despite these advances over the past years, insights into the landscape of structural variation remains lacking in many species of biomedical, behavioral, and evolutionary interest.

Native to the neotropical forests of Brazil and Peru, diurnal coppery titi monkeys (*Plecturocebus cupreus*; formerly classified as *Callicebus cupreus*; Groves 2005) form family groups consisting of a socially monogamous (bonded) pair and their offspring that often span multiple generations (Kinzey 1997). Females typically give birth to a single offspring once per year; similar to other primates, coppery titi monkeys have altricial young, though a defining feature of their social system is the extensive involvement of fathers in infant care (Mendoza and Mason 1986; Valeggia et al. 1999). The combination of social monogamy and paternal care — two biological characteristics uncommon amongst non-human primates and other mammals (Lukas and Clutton-Brock 2013) — make coppery titi monkeys a particularly valuable model system for studying neurobiology and social behavior (Bales et al. 2007, 2021; and see the review of Bales et al. 2017). For example, coppery titi monkeys have been used to study the effects of intranasally administered oxytocin — a neurohormone that plays a crucial role in pair bonding as well as in a variety of other social behaviors (Carter et al. 2020; Bales et al. 2021; Rigney et al. 2022) — and social interactions (Arias-del Razo et al. 2022a,b; Zablocki-Thomas et al. 2023; Witczak et al. 2024), with additional emerging evidence from clinical studies in humans suggesting that intranasally administered oxytocin might be a potential treatment to reduce social impairment in individuals affected by autism spectrum disorder (e.g., Moerkerke et al. 2024; and see the review of Horta et al. 2020). Yet, despite their importance in behavioral research, current knowledge regarding the underlying population genomic variation of coppery titi monkeys remains limited.

Combining novel high-coverage sequencing data from 26 coppery titi monkeys with a previously established, highly accurate multi-algorithm ensemble approach, this study presents the first map of structural variation in the species. Although the majority of identified variants were located in intergenic regions of the genome, several were predicted to be of high importance functionally. Amongst these variants, six impacted genes associated with human disease, including an inversion predicted to affect the PINK1-PARKIN pathway which has previously been linked to early-onset Parkinson’s disease. Taken together, these first insights into the structural variant landscape provided here will thus not only improve the usage of coppery titi monkeys as a behavioral model system, but also provide new avenues for future investigations of the biological mechanisms underlying human biology, health and disease.

## MATERIALS AND METHODS

### Animal subjects

This study was performed in compliance with all regulations regarding the care and use of captive primates, including the NIH Guidelines for the Care and Use of Animals and the American Society of Primatologists’ Guidelines for the Ethical Treatment of Nonhuman Primates. Procedures were approved by the UC-Davis Institutional Animal Care and Use Committee (protocol 22523).

### Samples, sequencing, and read mapping

Blood samples of 26 captive coppery titi monkeys (12 females and 14 males) were obtained from the pedigreed colony housed at the California National Primate Research Center. For each sample, genomic DNA was isolated, a PCR-free library was prepared to guard against amplification errors, and the whole genome was sequenced (Illumina NovaSeq 6000; 150 bp × 150 bp) to >50-fold coverage on average. The coverage in this study is thus considerably higher than that generally recommended (20×; Wold et al. 2021), with higher levels of coverage shown to increase both the accuracy and sensitivity of structural variant detection (Ahmad et al. 2023). The resulting sequencing reads were quality-controlled using TrimGalore v.0.6.10 (https://github.com/FelixKrueger/TrimGalore) which trims paired-end reads in a synchronized manner, discarding any pairs where at least one of the reads exhibits a length of < 20 bp after trimming Illumina adapter sequences and bases with a quality score < 20 from the 3’-ends. The pre-processed paired-end reads were aligned to the species-specific reference genome (PleCup_hybrid; GenBank accession number: GCA_040437455.1; Pfeifer et al. 2024) using BWA-MEM v.0.7.17 (Li 2013) and the resulting read alignments were sorted by coordinate using SAMtools *sort* v.1.16 (Danecek et al. 2021). Alignments from different runs of the same sample were merged using the Genome Analysis Toolkit (GATK) *MergeSamFiles* v.4.2.6.1 (van der Auwera and O’Connor 2020); subsequently, optical duplicates were flagged using *MarkDuplicates* in order to avoid potential biases in variant calling arising from artificially inflated read coverage (Pfeifer 2017). As high error rates, non-uniform coverage, and irregular insert size distributions can impede the discovery of structural variants (Mahmoud et al. 2019), SAMtools v.1.16 (Danecek et al. 2021) and goleft v.0.2.6 (https://github.com/brentp/goleft) were used to evaluate the quality and coverage patterns of the duplicated-marked mappings, respectively. Due to the higher quality and depth of coverage, analyses presented in this study focused on the autosomes (i.e., chromosomes 1-22). Sample information and mapping statistics are provided in Supplementary Table 1; and see Supplementary Figure 1 for the read coverage across autosomes.

### Structural variant discovery

Structural variants were discovered via an ensemble approach consisting of three short-read callers: DELLY (Rausch et al. 2012), Lumpy (Layer et al. 2014), and Manta (Chen et al. 2016). First, integrating paired-end mapping, split-read, and read depth signals from the read alignments, structural variants were jointly called in the study cohort using DELLY *call* v.1.2.6. Second, relying on the same signals as DELLY, in Lumpy v.0.2.13 (executed via Smoove *call* v.0.2.6; https://github.com/brentp/smoove) breakpoint likelihoods were modelled using a probabilistic framework to jointly call structural variants in the cohort. Based on the evidence observed at these breakpoints, the Bayesian likelihood method SVTyper v.0.7.0 (Chiang et al. 2015) was then used to determine the genotype likelihood for each sample. To limit the number of false positives in the dataset, and following the developers’ guidelines: (1) variants with a mean Smoove heterozygote quality (MSHQ) score ≤ 3 were excluded to remove heterozygous sites with a low genotyping confidence, (2) deletions with a < 0.7 fold-change in coverage relative to flanking regions (DHFFC) were filtered out to guard against alignment errors (particularly in regions of low complexity), and (3) duplications with a > 1.3 fold-change in coverage relative to genomic regions with similar GC-content (DHBFC) were removed to guard against artefacts resulting from GC-bias. Third, Manta v.1.6.0 was used to jointly call structural variants in the cohort by initially determining rough candidate breakpoint regions based on paired-end mapping and split-read signals only and subsequently performing a local *de novo* assembly in these candidate regions to resolve the breakpoints at base-pair resolution.

For each tool-specific call set, variants that passed the internal quality criteria were filtered following Wold et al. 2025, excluding any deletions shorter than 50 bp, duplications shorter than 300 bp, and inversions shorter than 300 bp given the difficulty of reliably distinguishing genuine signals from sequencing and alignment errors with short-read data at these scales; additionally, any variants longer than 50 kb were excluded to guard against unresolved repeats and other reference assembly issues. Lastly, tool-specific call sets were merged using SURVIVOR *merge* v.1.0.7 (Jeffares et al. 2017) to obtain consensus calls of structural variants detected by at least two of the three callers (requiring a 500 bp maximum distance between breakpoints of identical type).

### Structural variant genotyping

The consensus call set was re-genotyped using Graphtyper2 *genotype_sv* v.2.7.2 (Eggertsson et al. 2019). Following the developers’ recommendations, the “aggregate” genotyping model was used for copy number variants and insertions, the “breakpoint” model for inversions, and segregating variants were limited to those for which each individual in the cohort passed all internal sample-level filters. For downstream analyses, the final dataset was separated into sites consistent with the patterns of Mendelian inheritance and those violating these patterns using the *mendelian* plugin in BCFtools v.1.10 (Danecek et al. 2021).

### Structural variant annotation

SnpEff v.5.2 (Cingolani et al. 2012) was used to predict the functional impact of the discovered structural variants (note that SnpEff does not provide annotations for insertions). Afterward, structural variants classified as high impact were limited to those exhibiting the highest confidence (following the guidelines for human data; for details see Eggertsson et al. 2019) and then manually curated to evaluate potential medical relevance. This assessment relied on information available in the database of Disease-Gene Associations with annotated Relationships among genes (eDGAR; Babbi et al. 2017), which integrates pathogenicity information from UniProt’s Humsavar database (UniProt Consortium et al. 2015), NIH’s ClinVar database (Landrum et al. 2016, 2018) and the Online Mendelian Inheritance in Man (OMIM) database (Amberger et al. 2017). Additionally, a gene enrichment analysis was performed using the DAVID web server (Sherman et al. 2022).

## RESULTS AND DISCUSSION

### Discovery of structural variants in the coppery titi monkey

Structural variants were discovered by sequencing the genomes of 26 coppery titi monkeys to a mean individual coverage of 51.2× (range: 39.6× to 75.1×; Supplementary Table 1) and applying an ensemble approach that combines three established variant callers (DELLY [Rausch et al. 2012], Lumpy [Layer et al. 2014], and Manta [Chen et al. 2016]) to maximize sensitivity and specificity. This strategy has been demonstrated to perform well in primates (Subramanian et al. 2024; and see the benchmarking studies of Kosugi et al. 2019 and Gabrielaite et al. 2021). To further improve accuracy, discovered variants were re-genotyped using Graphtyper2 (Eggertsson et al. 2019), a population-scale genotyper that has been shown to improve genotype consistency across cohorts using pangenome graphs. Using this approach, a set of 13,492 structural variants were discovered among the 26 individuals analyzed in this study, with a mean of 3,042 variants per individual (range: 2,560–3,340; excluding the individual on which the reference assembly was based upon; Supplementary Table 2), spanning a combined 14.7 Mb of the autosomal genome. This diversity of structural variation is comparable to that observed in rhesus macaques (3,646 structural variants per individual; Thomas et al. 2021) but higher than that observed in humans from short-read data (∼2,100 to 2,500 structural variants per individual among the 2,504 individuals from the 26 populations included in the 1000 Genomes Project; Sudmant et al. 2015) as expected given the larger effective population size of coppery titi monkeys compared to humans (Terbot et al. 2026). It should be noted, however, that short-read based studies are unable to capture the full spectrum of structural variation and that more recent studies based on long-read data have been able to identify a much larger number of structural variants in the human genome (∼100,000 structural variants among 15 individuals; Audano et al. 2019). Consequently, the number of variants reported here certainly represents an underestimate and future work using cutting-edge long-read technologies will be required to provide a more comprehensive picture of the landscape of structural variation in the species, particularly with regards to large and complex events located within highly repetitive regions for which short-read based approaches have low recall rates (Kosugi and Terao 2024).

### Characterization of the landscape of structural variation in the coppery titi monkey

The vast majority of variants were copy number variants (92.8%; 12,484 deletions and 37 duplications), though a small number of unbalanced (765 insertions) and balanced structural variants (206 inversions) were also observed (see Figure 1a for genomic locations). Consistent with previous work showing that short-read approaches are biased towards the discovery of deletions (Pang et al. 2010), more deletions than duplications and insertions were detected in the coppery titi monkey genome. Although the ratio of deletions to duplications and insertions (∼16:1) was significantly higher than those observed in humans (*χ*2 = 13.303; df = 1; *p*-value = 0.000265) and other haplorrhines (rhesus macaques: *χ*2 = 19.438; df = 1; *p*-value = 1.04 × 10^-5^; Thomas et al. 2021), it was similar to that previously reported in strepsirrhines (aye-ayes: *χ*2 = 1.7751; df = 1; *p*-value = 0.1827; Versoza et al. 2025). In addition to genuine biological variation between the species, differences in both the available species-specific genomic resources and study design likely contribute to the observed differences. First, duplications tend to be poorly resolved in unfinished reference genome assemblies (Hartasánchez et al. 2018); additionally, recent work has shown that duplications and insertions are more difficult to detect than deletions using short-read approaches, with recall rates being inversely correlated to the size of these structural variants (Kosugi and Terao 2024). Consequently, differences in the quality of species-specific reference genomes could potentially lead to an underestimation of duplications in the coppery titi monkey and aye-aye genomes compared to human and rhesus macaque for which complete, or near complete, telomere-to-telomere assemblies are now available (Nurk et al. 2022; Zhang et al. 2025). Second, in contrast to the multi-algorithm ensemble approach applied in the studies of aye-ayes (Versoza et al. 2025) and coppery titi monkeys, the short-read variant catalogs previously generated for humans and rhesus macaques (Thomas et al. 2021) were based on a single caller (Lumpy) which may have missed genuine structural variation present in the genomes of these species (with recent benchmarking demonstrating a mean recall rate of only ∼3.5% using Lumpy; see Supplementary Table 10 in Kosugi et al. 2019).

**Figure 1.**
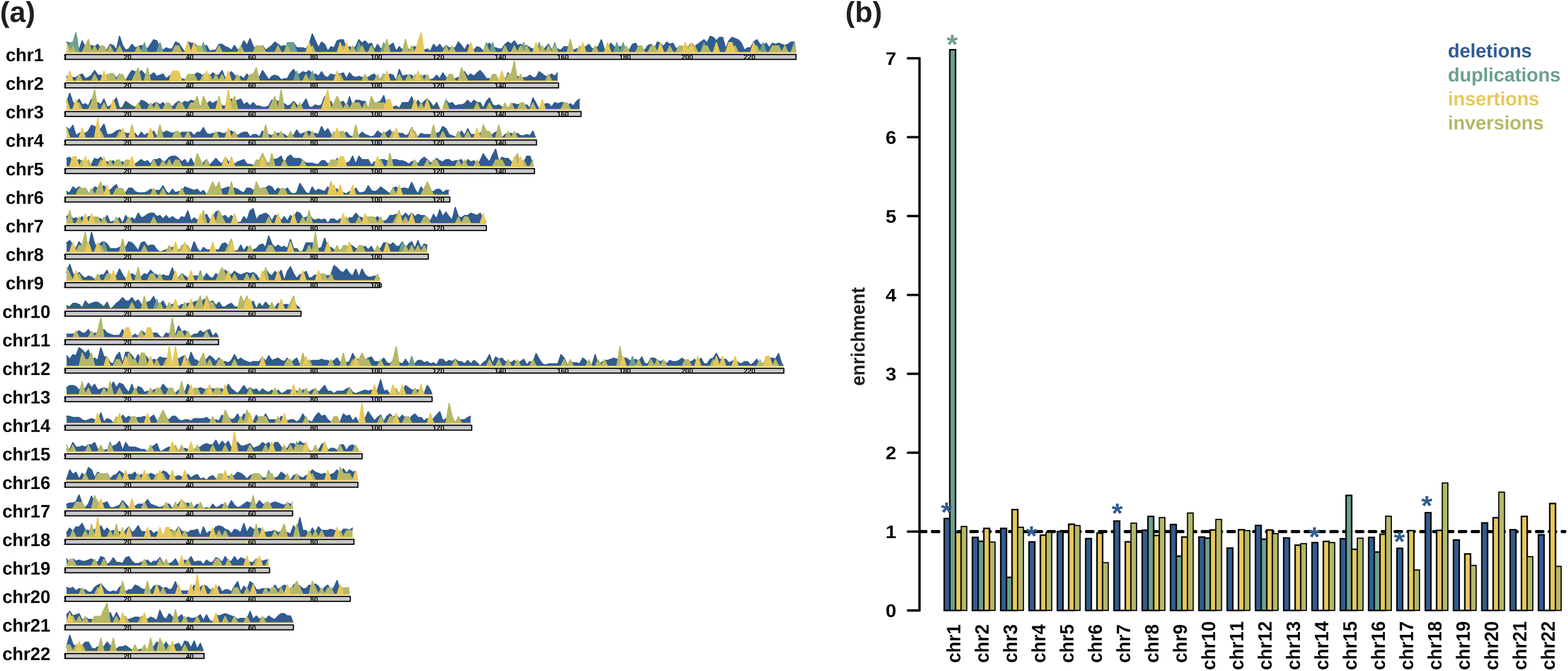
Landscape of structural variation in the coppery titi monkey genome. (a) Map of autosomal copy number variants (with deletions color-in blue and duplications in teal), insertions (yellow) and inversions (olive green). Chromosomes are displayed as horizontal bars and variants ividual peaks, with the height of each peak being proportional to the region length. (b) Distribution of structural variants across the autosomes. icant enrichments/depletions are marked by a star (*).

Similar to other species (Thomas et al. 2021; Versoza et al. 2025), the number of structural variants was strongly correlated with chromosomal length (deletions: *r* = 0.975, *p*-value = 1.33 × 10^-14^; duplications: *r* = 0.588, *p*-value = 4.03 × 10^-3^; insertions: *r* = 0.959, *p*-value = 1.96 × 10^-12^; inversions: *r* = 0.900, *p*-value = 1.22 × 10^-8^; Supplementary Figure 2). Although insertions and inversions were relatively evenly distributed between autosomes (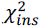 = 13.349, df = 21, *p*-value = 0.8959 and 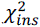 = 12.032, df = 21, *p*-value = 0.9388), deletions were significantly enriched on chromosomes 1 (1.17-fold enrichment; FDR-adjusted *p*-value = 1.16 × 10^-6^), 7 (1.13-fold; FDR-adjusted *p*-value = 8.17 × 10^-3^), and 18 (1.24-fold; FDR-adjusted *p*-value = 1.06 × 10^-5^) and depleted on chromosomes 4 (0.87-fold; FDR-adjusted *p*-value = 6.26 × 10^-3^), 14 (0.86-fold; FDR-adjusted *p*-value = 6.26 × 10^-3^), and 17 (0.79-fold; FDR-adjusted *p*-value = 1.31 × 10^-3^) (Figure 1b). Duplications were also significantly enriched on chromosome 1 (7.11-fold; FDR-adjusted *p*-value = 2.74 × 10^-27^); however, this observation should be interpreted with some caution given the small number of duplications in the dataset (*n* = 37). Moreover, structural variants were not randomly distributed, instead their genome-wide distribution was elevated in telomeric and sub-telomeric regions (Figure 1a) which tend to experience higher rates of non-allelic homologous recombination known to facilitate structural variation, particularly near transposable elements and other repetitive regions (Young et al. 2020).

Consistent with previous work in primates (Brandler et al. 2016; Brasó-Vives et al. 2020; Thomas et al. 2021; Subramanian et al. 2024; Versoza et al. 2025), both insertions and deletions were relatively short (with median lengths of 61 bp and 309 bp, respectively; Figure 2a) — an observation which likely reflects a combination of both biological constraints and technological limitations. On the one hand, errors during DNA replication and double-strand repair frequently generate short insertions and deletions; additionally, as longer insertions and deletions are more prone to disrupt protein-coding or regulatory regions, cause frameshifts, or change chromatin structure, they are more likely to be deleterious (Taylor et al. 2004; Itsara et al. 2010; Mills et al. 2011; Yang et al. 2024) and are thus expected to be purged from the population via purifying selection. On the other hand, although many short-read structural variant callers perform well in detecting short insertions and deletions, larger events as well as duplications and insertions are inherently more challenging to reliably identify, often leading to an ascertainment bias (Conrad and Hurles 2007; Sudmant et al. 2015; Kosugi et al. 2019; Mahmoud et al. 2019; Delage et al. 2020). In contrast, duplications and inversions typically impacted a greater number of nucleotides per event (with median lengths of 1.2 kb and 2.2 kb, respectively; Figure 2b).

**Figure 2.**
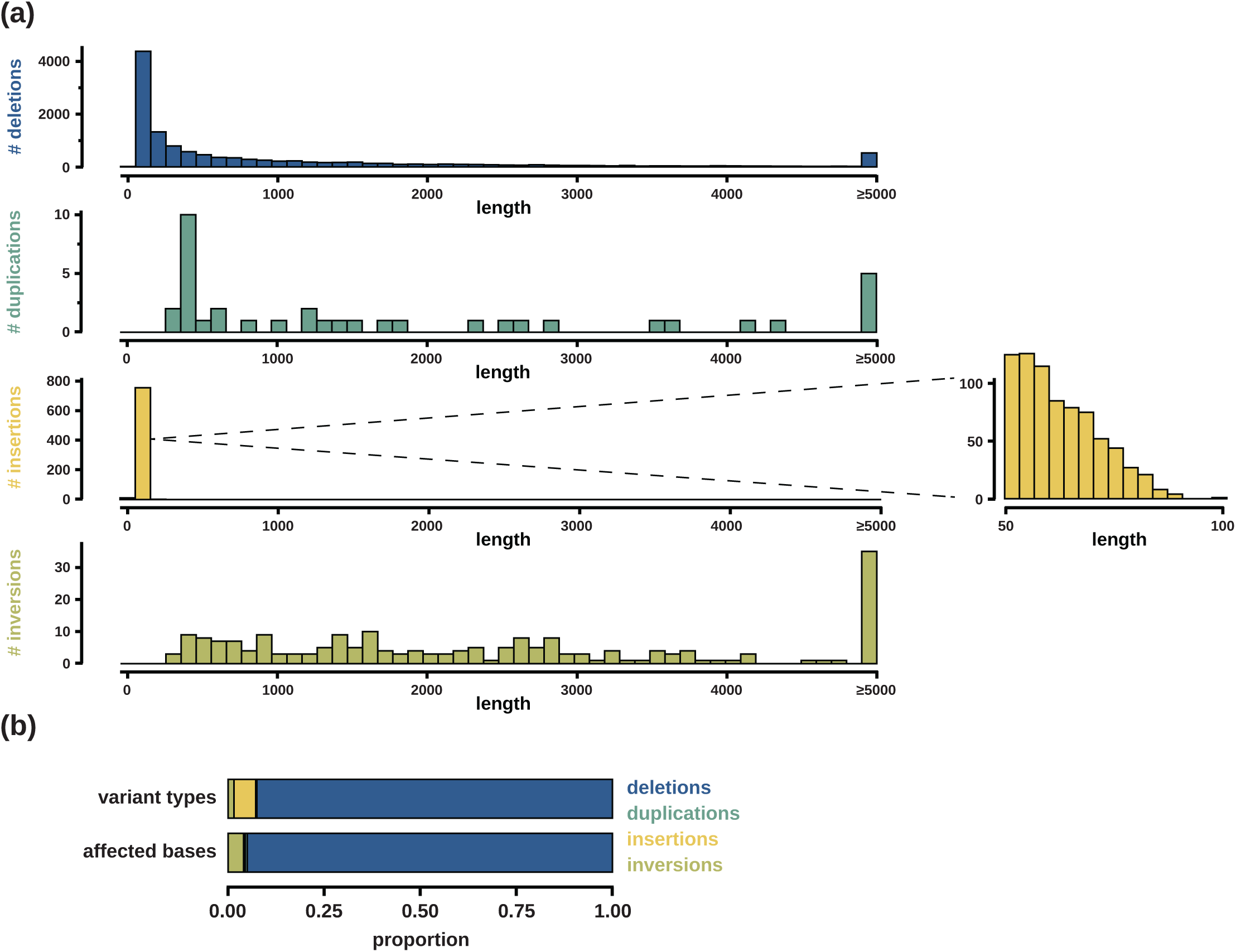
Characteristics of structural variation in the coppery titi monkey genome. (a) Length distribution of autosomal copy number variation (with deletions color-coded in blue and duplications in teal), insertions (yellow) and inversions (olive green). (b) Proportion of the base-pairs affected by the different variant types.

### Functional role of structural variants in the coppery titi monkey

In order to predict functional importance, structural variants were intersected with the annotations of the reference genome (Pfeifer et al. 2024) using SnpEff (Cingolani et al. 2012) to identify those impacting protein-coding genes, and the potential medical relevance of large-effect variants was subsequently evaluated using the human database of gene-disease associations, eDGAR (Babbi et al. 2017). The majority of variants was located within intergenic regions (65.08%; Supplementary Table 3), a significant enrichment compared to the overall genome composition (*χ*2 = 386,673; df = 6; *p*-value < 2.2 × 10^-16^). Although these variants are expected to be of limited functional importance — and consequently are frequently classified as modifiers or variants of low to moderate effect (Figure 3) — they can nevertheless have profound impacts on gene regulatory landscapes, for example, by altering enhancer / promotor functions or changing the boundaries of topologically associated domains. In fact, recent research has shown that structural variants harbored in intergenic regions frequently experience moderately strong purifying selection, suggesting that they are often detrimental to fitness (Saxena and Baer 2025). Additional modifiers included splice site variants (0.71%) as well as those located within transcripts (0.27%), 3’-UTRs (0.01%) and 5’-UTRs (0.02%). Of the variants residing within genic regions, many were predicted to be of high functional impact. Thereby, structural variants with predicted major effects were significantly enriched in cytoplasmic and membrane-associated cellular components, many of which localized to extracellular exosomes (73.2-fold enrichment; Supplementary Table 4). At the molecular level, they were characterized by enzyme-binding activities (52.7-fold enrichment), with a notable overrepresentation of GTPase activity, GTP-binding, and RNA-binding, consistent with roles in cell signaling, protein transport, and RNA regulation (Wittinghofer and Vetter 2011). Among the structural variants with predicted major effects, six were found to be located within disease-linked genes (Table 1).

**Figure 3.**
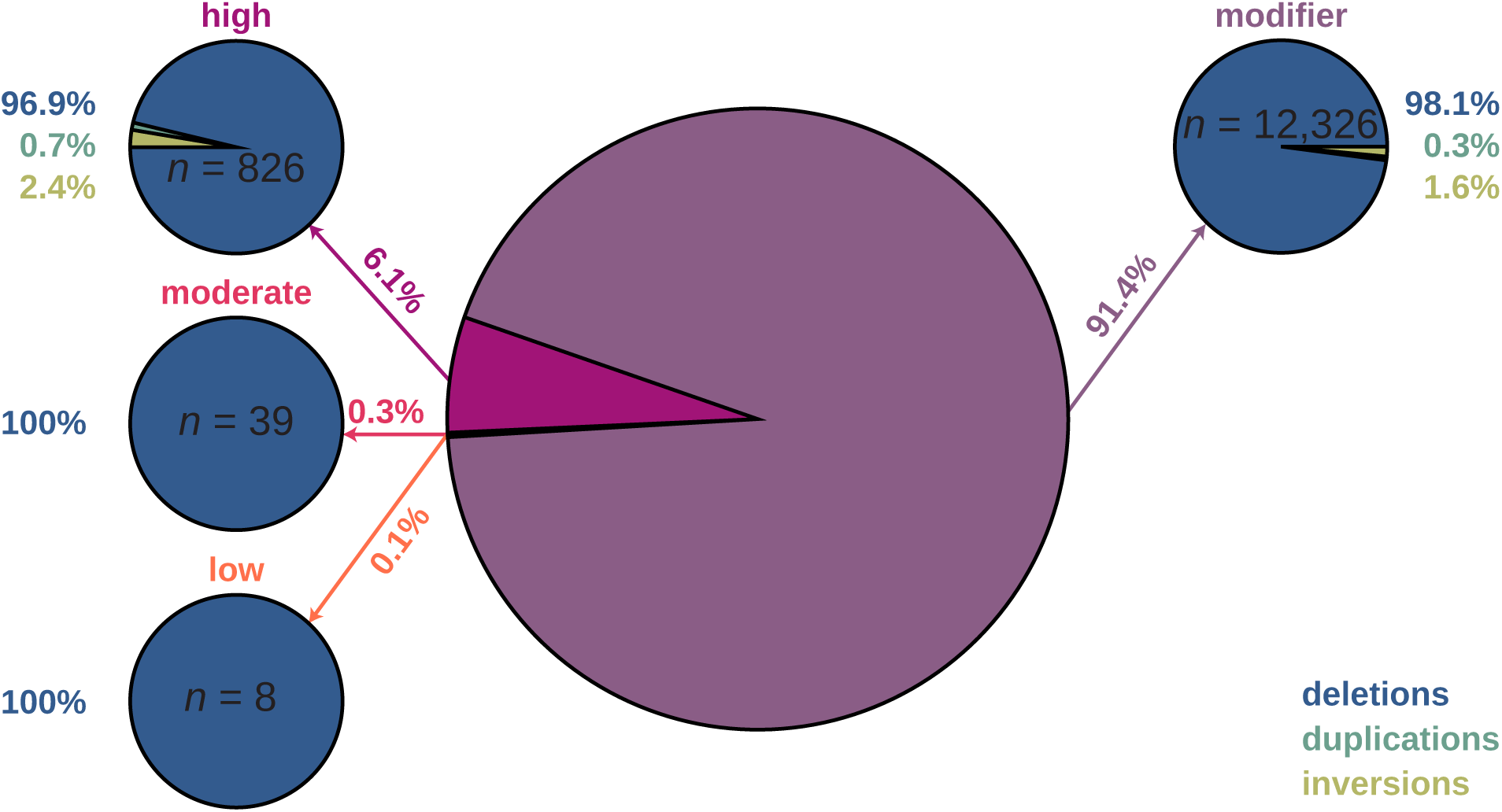
Functional annotation of structural variation in the coppery titi monkey genome. The proportion of autosomal copy number variation (with deletions color-coded in blue and duplications in teal), insertions (yellow) and inversions (olive green) predicted within non-coding or regulatory regions (i.e., modifiers; shown in purple) as well as those predicted to be likely disruptive (high: pink), more subtly affect protein function or gene regulation (moderate: rose), and unlikely to have a major functional impact (low: orange). Note that due to their size, several structural variants were predicted to have multiple effects.

**Table 1.** Structural variants with major effects predicted to affect disease-linked genes.

| type | chr | start | size | predicted effect | allele freq. | human gene | human disease phenotype |
| --- | --- | --- | --- | --- | --- | --- | --- |
| DEL | 1 | 10,718,272 | 1,041 | exon loss variant<br>splice acceptor/donor variant | 0.25 | ATL3 | hereditary sensory & autonomic neuropathy |
|  | 1 | 136,329,129 | 1,569 | frameshift variant<br>splice acceptor variant | 0.42 | ANKRD26 | thrombocytopenia |
|  | 1 | 167,808,572 | 68 | frameshift variant<br>start lost | 0.15 | PKP2 | arrhythmogenic cardiomyopathy |
|  | 1 | 183,000,971 | 534 | frameshift variant<br>splice donor variant | 0.06 | SALL4 | IVIC syndrome<br>Duane-radial ray syndrome |
| INV | 1 | 201,462,663 | 7,384 | splice acceptor variant | 0.12 | AMACR | AMACR deficiency<br>congenital bile acid synthesis defect |
|  | 13 | 99,538,214 | 1,950 | splice acceptor/donor variant | 0.04 | PINK1 | Parkinson's disease (early-onset) |

A ∼2.0 kb inversion was predicted to result in a change of a splice site in PTEN-induced putative kinase 1 (PINK1) which senses mitochondrial damage during cellular stress and that, depending on the severity of the damage, initiates either mitochondrial biogenesis or the removal of dysfunctional mitochondria via mitophagy (O’Callaghan et al. 2023). In humans, mutations in PINK1 have been linked to early-onset Parkinson’s disease — a neurodegenerative condition typically characterized by a variety of motor symptoms (such as bradykinesia, rigidity, and tremor) resulting from progressive neuronal cell death (Albanese et al. 2005). Parkinson’s disease has an estimated prevalence of 1% in individuals over the age of 65 years (Emborg 2017). Both pathogenicity and zygosity of mutations are known to impact the severity of the disease phenotype; the condition generally manifests around the age of 35 years in homozygous individuals and around the age of 43 years in heterozygous individuals, with females displaying an earlier onset than males (see the review by Yin and Dieriks 2025). Due to their close evolutionary relatedness and similar neuroanatomy, behavior, cognitive function, development, motor skills and aging process, non-human primates (including common marmosets, vervet monkeys, cynomolgus and rhesus macaques) are important models to study the environmental and genetic risk factors underlying the disease and to develop neuroprotective and treatment strategies such as dopamine replacement and gene therapies (see the reviews of Emborg 2017; Pan et al. 2024, and references therein). For example, several recent studies used CRISPR/Cas9-mediated knockouts of PINK1 to demonstrate that the disruption of this gene leads to neuronal loss in the cerebral cortex and Parkinson-like symptoms in non-human primates (Yang et al. 2019, 2022; Li et al. 2020a, 2021). Although such studies are frequently based on transgenic or chemically-induced models, recent research suggests that aged non-human primates can also naturally develop symptoms resembling those observed in humans affected by the disease (Hurley et al. 2011; Li et al. 2020a, 2021).

A ∼7.4 kb inversion was predicted to result in a change of the splice acceptor site in alpha-methylacyl-CoA racemase (AMACR), an enzyme aiding the metabolism of fatty acids and the synthesis of bile acid during digestion. Mutations in this gene have been found to lead to AMACR deficiency in humans which has been linked to a variety of adult-onset neurodegenerative issues ranging from migraines to visual impairments, neuropathy, cognitive decline, and seizures (Ferdinandusse et al. 2000; Thompson et al. 2008; Smith et al. 2010; Dick et al. 2011; and see the review of Clarke et al. 2004). Additionally, there has been evidence that a homozygous mutation in AMACR can cause congenital bile acid synthesis defect type 4, a rare genetic disorder characterized by cholestatic liver disease (Ferdinandusse et al. 2000; Setchell et al. 2003).

A 68 bp deletion was predicted to result in the loss of a start codon through a frameshift in the coding region of plakophilin-2 (PKP2) — an integral component of both cardiac desmosomes, cell-cell adhesion complexes that provide both structural and mechanical integrity to the cardiac muscle (Mertens et al. 1996), and the cardiac connexome (Cerrone et al. 2017). In humans, multiple deletions, insertions, non-sense mutations, missense mutations, as well as splice site mutations in this gene are known to cause arrhythmogenic cardiomyopathy — a heart disease with an estimated prevalence of 1 in 1,000 to 5,000 individuals (Basso et al. 2009) that is characterized by defects in cardiac morphogenesis through a fibrofatty replacement of the myocardium, and that is associated with cardiac arrhythmias and sudden cardiac arrest (e.g., Thiene et al. 1988; Gerull et al. 2004; Dalal et al. 2006; van Tintelen et al. 2006; Kirchner et al. 2012; and see the review of Corrado et al. 2017).

A ∼0.5 kb deletion was predicted to result in a frameshift in the spalt like transcription factor 4 (SALL4). In humans, mutations in this gene have been implicated in two disorders with similar phenotypes: IVIC syndrome (Paradisi and Arias 2007) and Duane-radial ray syndrome (also known as Okihiro syndrome; Al-Baradie et al. 2002; Kohlhase et al. 2002, 2003; Borozdin et al. 2004; Miertus et al. 2006), both characterized by abnormalities of the upper limbs, motoneuron development as well as a range of other features, often with variable expressivity.

A 1.0 kb deletion was predicted to lead to the loss of an exon in the membrane-anchored GTPase Atlastin-3 (ATL3) gene. Missense mutations in this gene are known to cause a heritable form of sensory neuropathy (hereditary sensory neuropathy type 1F) in humans which can lead to chronic ulcerations, osteomyelitis, and acro-osteolysis of the lower limbs in aging affected individuals (Fischer et al. 2014; Kornak et al. 2014; Xu et al. 2019).

Lastly, a 1.6 kb deletion was predicted to result in a frameshift in the ankyrin repeat domain-containing protein 26 (ANKRD26). In humans, there is evidence that mutations in this gene can cause thrombocytopenia-2 — a condition characterized by an abnormally low number of thrombocytes in the blood of the affected individuals which often causes relatively mild symptoms, including easy bruising and a minor bleeding tendency (Savoia et al. 1999; Drachman et al. 2000).

### *De novo* structural variants in the coppery titi monkey

In many species, *de novo* structural variants occur at orders of magnitude lower frequency than point mutations (Belyeu et al. 2021); for example, in humans, high-resolution studies of large cohorts of families observed one *de novo* copy number variant per 3.5 births (with a projected rate of one *de novo* mutation per 2-8 births; Collins et al. 2020). A total of 10 *de novo* structural variants — eight deletions, one duplication, and one insertion — were identified among the parent-offspring trios included in this study, i.e., one in every 1.5 births. This rate is similar to the rate of one in every 1.75 births previously reported in rhesus macaques (i.e., seven deletions and one duplication among 14 trios; Thomas et al. 2021). The higher rate of *de novo* structural mutations observed in coppery titi monkeys compared to those observed in rhesus macaques and humans might, at least in part, be driven by differences in generation time (with species exhibiting shorter generation times expected to experience higher rates of molecular evolution; Ohta 1993). As anticipated from the overall landscape of structural variation, the majority of the *de novo* variants (60%) were located within intergenic regions (Supplementary Table 5); the single exonic variant — a 570 bp deletion — was located within a gene of unknown function (KAL0613339).

## CONCLUSION

As a primate of considerable biomedical and behavioral interest, gaining novel insights into the heritable variation characterizing the genome of the coppery titi monkey is crucially important to advance both on-going and future research endeavors related to human health and disease. Moreover, in contrast to haplorrhines, population-scale structural variant catalogs remain scarce in most platyrrhines and thus, this first map of the genomic architecture of structural variation for a representative of the Pitheciidae family will serve as a valuable genomic resource for future evolutionary studies across the primate clade. Nevertheless, even though the high-coverage data presented here offers a first glimpse into the structural variant landscape of the species, future long-read studies will be needed both to characterize the full spectrum of genomic variation — particularly with regards to insertions and inversions as well as large and complex events located within repetitive regions which are systematically under-detected using short-read data — as well as to improve breakpoint resolution, particularly for inversions which are frequently flanked by segmental duplications of near-perfect sequence identity (Porubsky and Eichler 2024). Helpfully, as the cost and sample requirements of long-read sequencing technologies continue to decrease, novel studies are expected to emerge that will allow for a more comprehensive understanding of the causes and consequences of structural variation across different primate lineages.

## ACKNOWLEDGEMENTS

DNA extraction, library preparation, and Illumina sequencing were conducted at the DNA Technologies and Expression Analysis Core at the UC Davis Genome Center (supported by NIH Shared Instrumentation Grant 1S10OD010786-01) and Novogene (Sacramento, CA, USA). Computations were performed on the Sol supercomputer at Arizona State University (Jennewein et al. 2023).

