## Supplementary Materials for "The landscape of structural variation in coppery titi monkeys (*Plecturocebus cupreus*)"

**Supplementary Table 1**. Sample information and mapping statistics.

| sex | individual | coverage | % mapped | % properly paired |
|---|---|---|---|---|
| male | I1 | 65.9 | 99.86 | 98.14 |
| | I2 | 45.2 | 99.92 | 98.78 |
| | I3 | 40.9 | 99.94 | 98.79 |
| | I4 | 39.6 | 99.87 | 98.82 |
| | I5 | 56.7 | 99.84 | 98.07 |
| | I6 | 62.8 | 99.88 | 98.19 |
| | I7 | 61.8 | 99.84 | 98.04 |
| | I8 | 48.9 | 99.85 | 97.95 |
| | I9 | 43.6 | 99.94 | 98.89 |
| | I10 | 52.0 | 99.86 | 97.92 |
| | I11 | 61.8 | 99.86 | 98.02 |
| | I12 | 55.7 | 99.85 | 97.86 |
| | I13 | 50.1 | 99.85 | 97.94 |
| | I14 | 46.3 | 99.90 | 98.64 |
| female | I15 | 42.2 | 99.89 | 98.74 |
| | I16 | 61.1 | 99.81 | 97.29 |
| | I17 | 46.0 | 99.92 | 98.77 |
| | I18 | 59.2 | 99.85 | 97.81 |
| | I19 | 47.2 | 99.84 | 97.87 |
| | I20 | 42.3 | 99.92 | 98.70 |
| | I21 | 41.0 | 99.90 | 98.61 |
| | I22 | 41.1 | 99.92 | 98.54 |
| | I23 | 57.7 | 99.86 | 97.98 |
| | I24 | 42.1 | 99.94 | 99.02 |
| | I25 | 45.4 | 99.94 | 98.90 |
| | I26 | 75.1 | 99.86 | 98.44 |

**Supplementary Table 2**. Number of structural variants discovered in each individual.

| sex | individual | # variants |
|---|---|---|
| male | I1 | 1,969 |
| | I2 | 3,004 |
| | I3 | 3,079 |
| | I4 | 2,560 |
| | I5 | 3,136 |
| | I6 | 3,226 |
| | I7 | 3,040 |
| | I8 | 3,070 |
| | I9 | 2,874 |
| | I10 | 3,245 |
| | I11 | 3,127 |
| | I12 | 3,134 |
| | I13 | 3,207 |
| | I14 | 2,999 |
| female | I15 | 3,175 |
| | I16 | 3,079 |
| | I17 | 3,099 |
| | I18 | 3,195 |
| | I19 | 3,068 |
| | I20 | 2,921 |
| | I21 | 2,861 |
| | I22 | 2,952 |
| | I23 | 3,340 |
| | I24 | 2,748 |
| | I25 | 2,854 |
| | I26 | 3,055 |

**Supplementary Table 3**. Predicted structural variant effects by region and type.

| region | % of variants |
|---|---|
| intergenic | 65.08 |
| intron | 12.38 |
| exon | 7.96 |
| downstream | 6.81 |
| upstream | 6.50 |
| splice site acceptor | 0.41 |
| gene | 0.27 |
| transcript | 0.27 |
| splice site donor | 0.24 |
| splice site region | 0.05 |
| 5'-UTR | 0.02 |
| 3'-UTR | 0.01 |

| type | % of variants |
|---|---|
| intergenic region | 57.73 |
| intron variant | 14.15 |
| downstream gene variant | 6.04 |
| upstream gene variant | 5.77 |
| exon loss variant | 5.23 |
| splice region variant | 3.34 |
| splice donor variant | 2.44 |
| splice acceptor variant | 2.13 |
| frameshift variant | 1.42 |
| conservative in-frame deletion | 0.38 |
| disruptive in-frame deletion | 0.21 |
| bidirectional gene fusion | 0.18 |
| stop lost | 0.18 |
| inversion | 0.16 |
| start lost | 0.16 |
| transcript ablation | 0.14 |
| stop gained | 0.10 |
| intragenic variant | 0.08 |
| gene fusion | 0.04 |
| 5'-UTR truncation | 0.04 |
| duplication | 0.03 |
| feature ablation | 0.02 |
| non-coding transcript variant | 0.02 |
| 3'-UTR truncation | 0.01 |
| 3'-UTR variant | 0.01 |
| 5'-UTR variant | 0.01 |
| exon region | 0.01 |

**Supplementary Table 4.** Gene enrichment of structural variants with predicted major effects.

| | term | fold enrichment | FDR |
|---|---|---|---|
| **cellular component** | GO:0070062~extracellular exosome | 73.2 | 0.00083 |
| | GO:0005829~cytosol | 3.5 | 1.00000 |
| | GO:0016020~membrane | 3.5 | 0.35456 |
| | GO:0005737~cytoplasm | 2.6 | 0.82706 |
| **molecular function** | GO:0019899~enzyme binding | 52.7 | 0.27840 |
| | GO:0003924~GTPase activity | 14.3 | 0.21479 |
| | GO:0042802~identical protein binding | 13.2 | 0.21479 |
| | GO:0005525~GTP binding | 11.9 | 0.21479 |
| | GO:0003723~RNA binding | 6.9 | 0.38448 |
| | GO:0005515~protein binding | 6.6 | 0.00075 |

**Supplementary Table 5**. *De novo* structural variation detected across parent-offspring trios. Location (chromosome [chr] and start position), variant type (deletion [DEL], duplication [DUP], and insertion [INS]), and genomic annotation are defined according to the species genome assembly (note that SnpEff does not provide annotations for insertions).

| chr | start | size | region | type |
|---|---|---|---|---|
| 1 | 1,472,628 | 516 | intergenic | DUP |
| 1 | 2,745,819 | 550 | intergenic | DEL |
| 1 | 59,742,399 | 2,295 | intergenic | DEL |
| 1 | 135,542,913 | 74 | intronic | DEL |
| 3 | 24,252,895 | 144 | intergenic | DEL |
| 4 | 40,724,117 | 51 | – | INS |
| 6 | 123,701,983 | 2,216 | intergenic | DEL |
| 8 | 36,319,872 | 570 | exonic | DEL |
| 8 | 115,277,432 | 5,669 | intergenic | DEL |
| 14 | 130,238,324 | 58 | upstream | DEL |

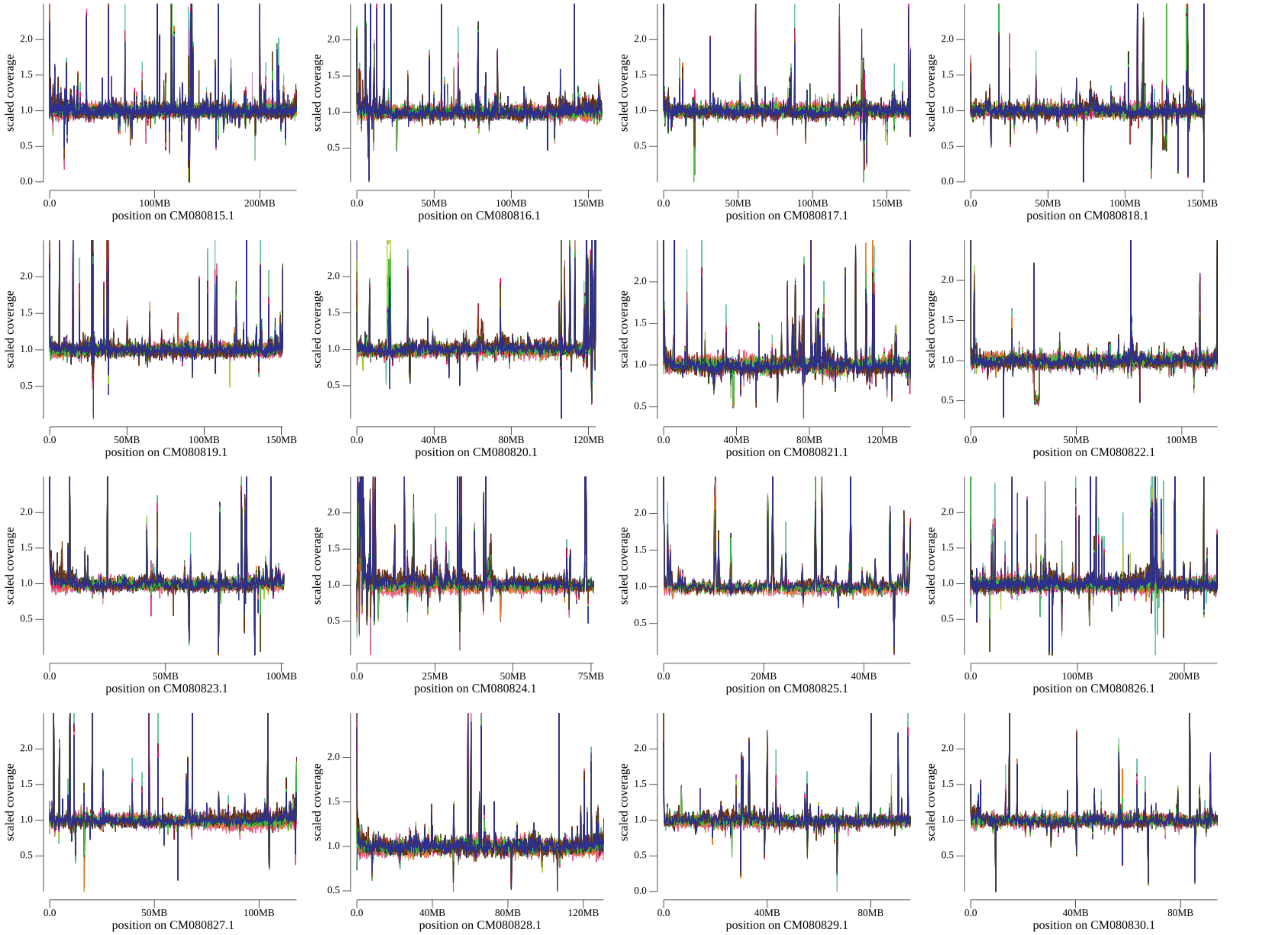
scaled coverage
position on CM080815.1
position on CM080816.1
position on CM080817.1
position on CM080818.1
position on CM080819.1
position on CM080820.1
position on CM080821.1
position on CM080822.1
position on CM080823.1
position on CM080824.1
position on CM080825.1
position on CM080826.1
position on CM080827.1
position on CM080828.1
position on CM080829.1
position on CM080830.1

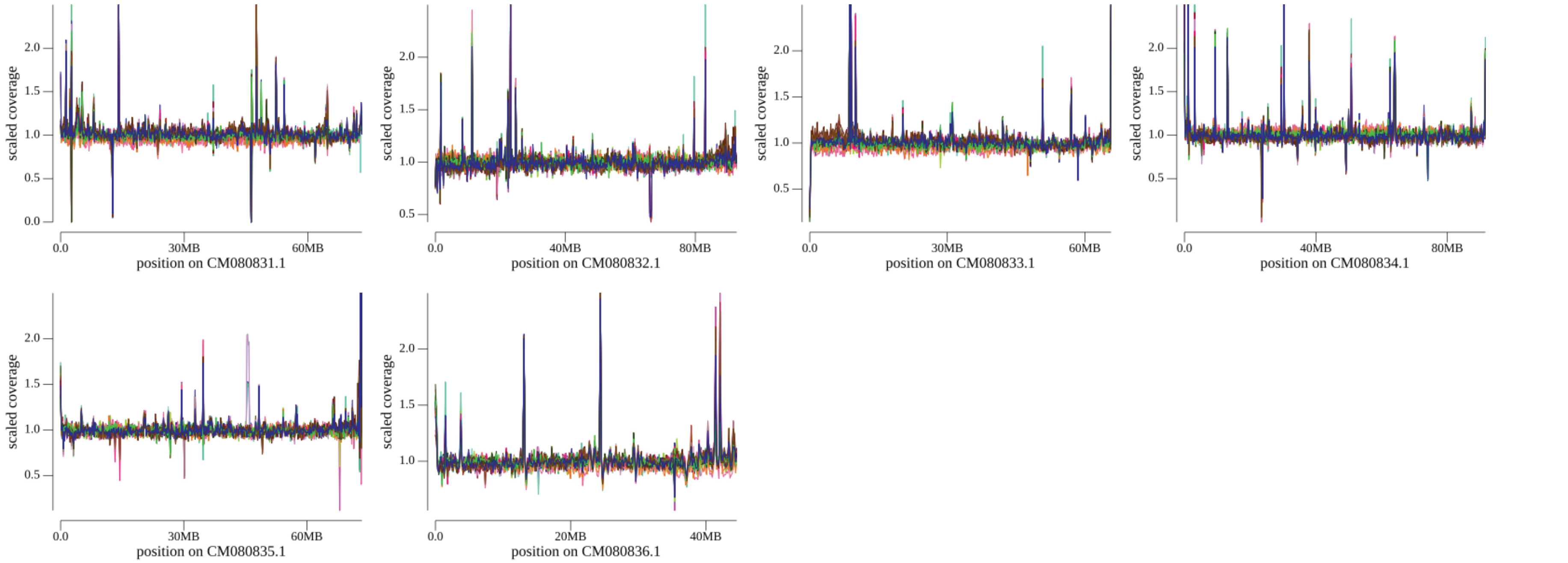


**Supplementary Figure 1**. Read coverage across autosomes.

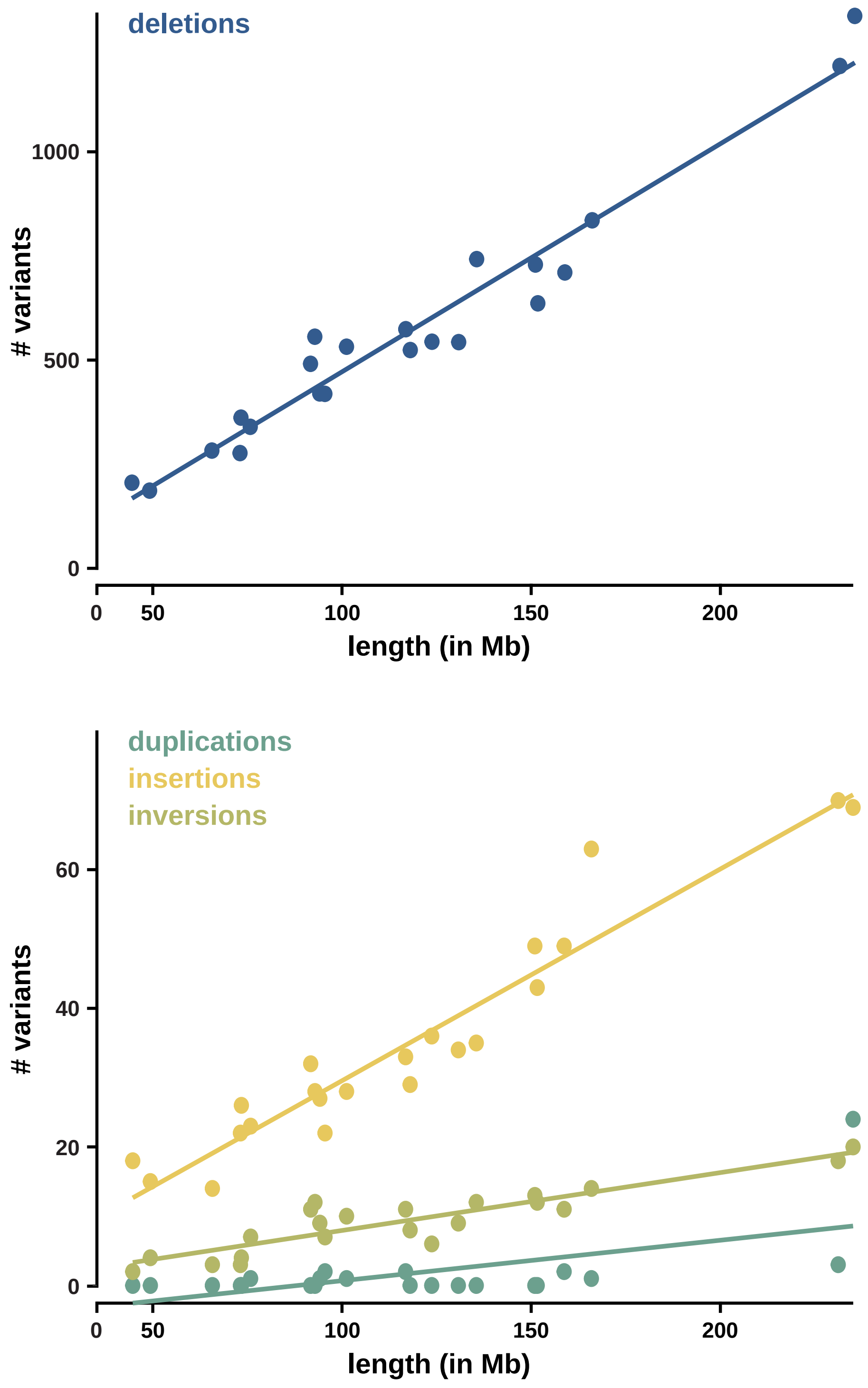


**Supplementary Figure 2**. Correlation between the number of deletions (color-coded in blue; top panel) and duplications, insertions, and inversions (in teal, yellow, and green, respectively; bottom panel) and autosomal length.
